# Isolation and Characterization of Extracellular Vesicles from Mouse Retina Tissue

**DOI:** 10.64898/2026.05.24.724732

**Authors:** Ishrat Ahmed, Olesia Gololobova, Mohammed Amanullah, Zach Troyer, Joshua H. Yu, James T. Handa, Jiang Qian, Seth Blackshaw, Kenneth Witwer

## Abstract

This protocol provides a standardized workflow for the isolation of extracellular vesicles (EVs) from mouse retinal tissue, and includes an assessment of EV size and concentration, marker expression, and EV visualization in accordance with the International Society for Extracellular Vesicles Minimal Information for Studies of Extracellular Vesicles (MISEV) guidelines. Most retinal EV studies rely on cell culture, which may not fully capture *in vivo* biology. Our approach more accurately reflects physiological and pathological EV states *in vivo* by enabling the extraction of EVs from intact retinal tissue. This method addresses a key gap in the field by providing a reproducible and rigorous protocol for studying retinal EVs in a biologically relevant context.

## Introduction

Extracellular vesicles (EVs) are membrane-bound particles released by cells that contain proteins, lipids, and nucleic acids. Increasing evidence suggests that EV cargo reflects the physiological state of the cell of origin and can contribute to disease processes, including photoreceptor degeneration, in the retina [1]. To date, most studies of retinal EVs have focused on vesicles isolated from cultured retinal pigment epithelium, photoreceptors, retinal progenitor cells, and Müller glial cells, or from biofluids such as aqueous and vitreous humor [2–8]. While these approaches have provided valuable insights, they may not fully capture the complexity of EV populations present within intact retinal tissue. Isolating EVs from small tissues such as the mouse retina presents inherent challenges due to limited starting material, which can restrict EV yield and limit downstream analyses. These constraints necessitate an optimized protocol to ensure sufficient recovery without compromising EV integrity or purity.

A wide variety of enrichment techniques are available to separate EVs, each exploiting different physical or biochemical properties. Common approaches include size-based methods such as differential ultracentrifugation, size exclusion chromatography, ultrafiltration, and sequential filtration [9, 10]. Additional strategies include density gradient centrifugation, immunoaffinity-based isolation, and precipitation-based methods. Combinations of these strategies are often used to improve purity or yield. Here, we describe a novel protocol for the isolation of mouse retinal EVs that combines differential centrifugation, size exclusion chromatography, and ultracentrifugation to improve EV yield and reproducibility.

Given the diversity of methods available for EV separation, it is essential to understand as much as possible about the EV product resulting from the chosen workflow. In its “Minimal Information for Studies of Extracellular Vesicles” (MISEV), the International Society for Extracellular Vesicles recommends: 1) measurement of EV concentration and size distribution using techniques such as nanoparticle tracking analysis, resistive pulse sensing, or nanoflow cytometry; 2) protein analysis through methods like Western blot, ELISA, flow cytometry, or single-particle interferometric reflectance imaging sensor (SP-IRIS); and 3) visualization, e.g., with transmission electron microscopy [11]. Our protocol emphasizes reproducibility and analytical rigor, consistent with these MISEV standards.

By enabling the direct extraction of EVs from intact retinal tissue, our approach captures EV populations that more accurately reflect physiological and pathological states *in vivo* relative to EVs obtained from *in vitro* models. This is particularly important for understanding intercellular communication within the retina and for identifying EV-associated biomarkers relevant to retinal diseases. This optimized protocol provides the necessary rigorous foundation for downstream applications, including molecular profiling and functional studies.

## Materials and Supplies

The names of the suppliers and catalog numbers for all materials and equipment used in retina dissection and EV separation are listed in **Table 1**. C57BL/6J mice (The Jackson laboratory, strain # 000664) were used for this protocol. All procedures involving mice are carried out in accordance with institutional and national animal care guidelines and have been approved by the Institutional Animal Care and Use Committee (IACUC) at Johns Hopkins University.

**Table 1.**
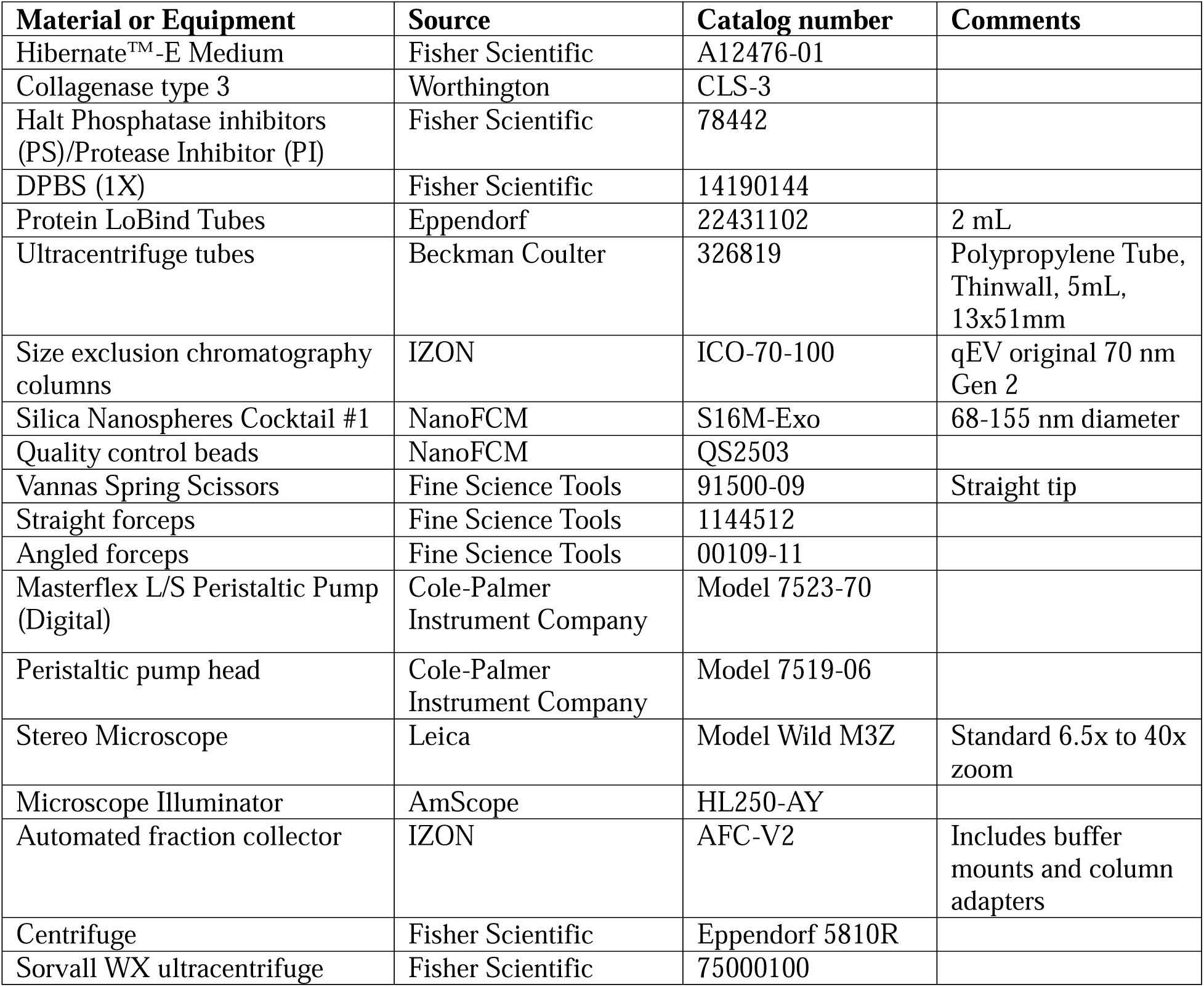
Names, sources and catalog numbers of materials and equipment required for retina dissection and EV separation.

### Detailed Methods

This section provides a detailed description of the procedures used for retinal dissection and EV separation, followed by EV analysis according to MISEV guidelines. The protocol is summarized in **Figure 1**.

**Figure 1.**
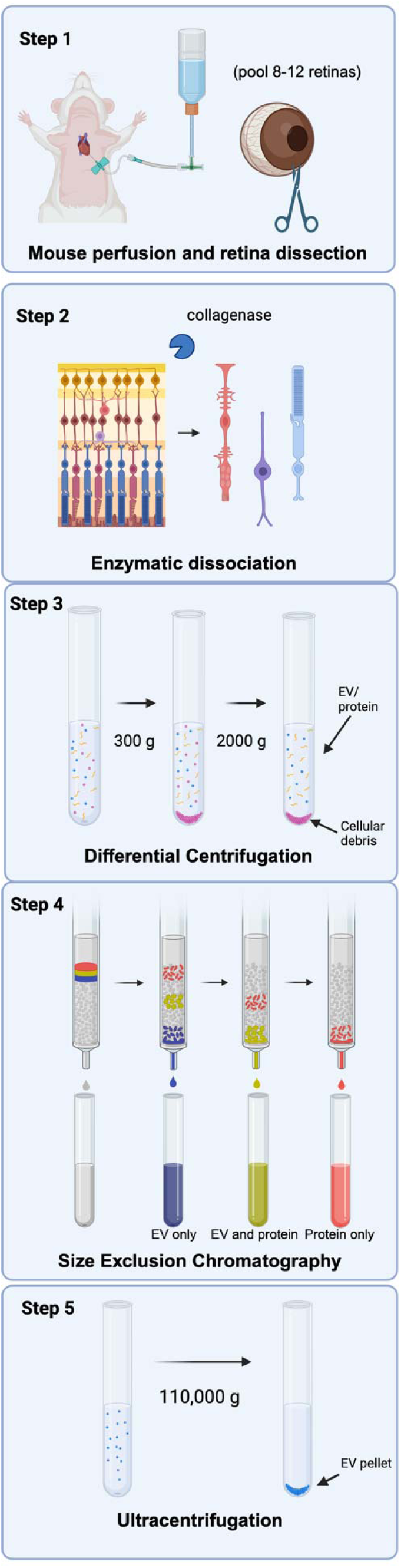
EVs are isolated from mouse retina using enzymatic dissociation, differential centrifugation, size exclusion chromatography, and ultracentrifugation followed by multi-modal characterization in accordance with MISEV guidelines.

#### 1. Preparation of stock solutions

The following stock solutions should be prepared in advance and can be stored at −20°C. Aliquot all stocks to avoid repeated freeze–thaw cycles.

1. **1x protease inhibitor (PI) / phosphatase inhibitor (PS) in D-PBS**: 2 tubes PI/PS in 19.8 mL D-PBS
2. **5x PI/PS in Hibernate®-E Medium**: 2 tube PI/PS in 3.8 mL Hibernate®-E Medium
3. **75 U/mL of collagenase in Hibernate®-E**:

3.75 mg 75 U/ml collagenase in 10 mL Hibernate®-E Medium (yields collagenase activity: 200 U/mg)
Prepare at least 400 uL of solution per 50 mg of tissue
Of note, it is essential to normalize by collagenase activity rather than weight to prevent inconsistent digestion between batches.

#### 2. Dissection of retinal tissue

Timing: 1-2 hours

During this step, we perfuse the mice, enucleate the eye, and dissect the retina for digestion. Rapid dissection of the retina followed by immediate freezing on dry ice is critical to preserve EV integrity, as it minimizes enzymatic activity and tissue degradation, which can alter EV composition and reduce sample quality.

1. Deeply anesthetize the mice and perfuse the mice with ice-cold PBS according to approved animal care protocols. This step is essential for removing blood-derived EVs.
2. Gently retract the eyelids using fine forceps and enucleate the eye.
3. Immediately place the enucleated eye on a cooled dissection plate with ice-cold PBS.
4. Puncture the cornea with a sterile needle. Insert Vannas micro-scissors into the hole and carefully cut along the limbus.
5. Excise the cornea and lens, leaving the posterior eye cup intact.
6. Remove the vitreous humor with fine forceps while avoiding traction on the retina.
7. Gently separate the neural retina from the surrounding retinal pigment epithelium by inserting angled forceps under the retina and bluntly dissecting.
8. Immediately transfer the tissue into a 2mL Protein LoBind microcentrifuge tube on dry ice. To obtain sufficient retinal tissue for EV isolation, we recommend pooling at least 8-12 mouse retinas.
9. Tissues may be stored at −80°C until used.

#### 3. Tissue preparation and dissociation

Timing: 30 minutes

1. Transfer the tissue from −80°C storage to dry ice. Keep the tissue on dry ice until indicated to maintain cellular and EV integrity.
2. Weigh the tissue (on dry ice)
3. Calculate the required volumes of buffers as follows: Tissue weight: **X** mg Volume of collagenase Hib-E buffer (stock solution #3): **V1** = 400ul * **X**/50 mg Volume of 5x PI/PS Hib-E buffer (stock solution #2): **V2** = **V1**/5
4. Add 400 µL per 50 mg of tissue of 75U/ml collagenase Hib-E solution (stock solution #3) as calculated in step 3 to the 2 mL tube and incubate for a total of 12 min at 37°C on a shaker at 350 rpm.

a. After 5 minutes of incubation, mix twice by inversion. Then, incubate for 5 more minutes.
b. Use a 1000 µL pipette tip with an enlarged opening (cut with a scalpel) to gently pipette up and down twice. It is essential to use an enlarged opening to limit shear stress.
c. Incubate for 2 min.
5. After incubation, place the tube containing digested tissue on ice immediately. Add ice-cold 5x Hib-E PI/PS solution (stock solution #2) as calculated in step 3 to the 2mL tube to quench the enzyme. Pipette only once with a 1000 µL pipette tip with an enlarged opening.

#### 4. Separation of retinal EVs

##### 4.1. Differential centrifugation for removal of cells, tissue debris, and large particles

Timing: 1 hr

1. Centrifuge digested tissue at 300 x g for 10 min at 4°C.
2. Remove the supernatant and transfer to a new tube.
3. Add 100 uL 1X PI/PS D-PBS (stock solution #1) to the pellet and store at −80°C (if needed later for protocol validation).
4. Centrifuge the supernatant at 2000 x g for 15 min at 4°C.
5. Transfer the supernatant to a new tube and record the volume. The supernatant is enriched for EVs. Keep on ice to prevent aggregation and decrease enzymatic activity.
6. Resuspend the residual pellet in 50 uL of 1X PI/PS D-PBS (stock solution #1) and store at −80°C (if needed later for protocol validation).

##### 4.2. Size exclusion chromatography for separation of EVs and proteins

Timing: 1-2 hours

1. Bring the volume of the 2000 x g supernatant (step 5 in section 4.1) to either 500 µL or 1000 µL with 1X PI/PS D-PBS buffer (stock solution #1). Use the recorded volumes of the supernatant to calculate the amount of buffer to add.
2. Set up the IZON automatic fraction collector. If using the qEV original columns, the following settings are recommended: fraction size 0.5 mL and void volume 2.4 mL. Fractions may also be collected manually or using another type of fraction collector.
3. Column preparation: Set up the column and wash with 17 mL (for 500 µL samples) or 27 mL (for 1000 µL samples) using room temperature PBS.
4. Transfer the sample to the column and press start.
5. Once the sample has fully entered the column and the column is no longer dripping, add either 8 mL of PBS or 14 mL PBS (as prompted by the instrument).
6. For a starting sample volume of 500 µL collect a total of 12 tubes. Fractions 1 to 4 contain EVs, fractions 5 to 8 contain mixed EVs and protein, and fractions 9 to 12 contain proteins. Each fraction will contain approximately 0.5 mL volume. For a starting sample volume of 1000 µL, 2 tubes are required to collect each fraction (for a total volume of 1 mL). Therefore, 24 tubes will be needed in total, and the first 8 tubes will contain EV-enriched fractions.

##### 4.3. Ultracentrifugation for EV concentration

Timing: 2 hrs

While this protocol uses ultracentrifugation to concentrate the EVs, other methods, such as ultrafiltration, may also be used.

1. Transfer the pooled EV-enriched fractions to an ultracentrifuge tube. Ensure that the tubes are nearly filled. If needed, mark 0.5 cm from the top and fill to the line with 5x Hib-E PI/PS buffer (stock solution #2). It is essential to fill ultracentrifuge tubes to ensure balance and prevent tube collapse.
2. Centrifuge for 70 minutes using a AH-650 pre-chilled rotor (capacity: 4.5 mL) at 110,000 x g (Average deceleration=9, acceleration=9) at 4°C.
3. Remove the tubes from the rotor immediately after completion. Remove one tube from the rotor at a time and transfer the supernatant by pouring. Repeat with the remaining tubes.
4. Resuspend pellets in 30 µl ice cold 1X PS/PI DPBS as follows. Pipette up and down 10 times, vortex for 30 seconds, and place on ice for 20 minutes. Then, pipette up and down 10 times and vortex for 30 seconds.
5. Transfer to 1.5 mL protein LoBind tubes and record the volume transferred (expected range 35-50 µl). Of note, when choosing a tube type and manufacturer, use particle-free buffer to assess possible shedding of nanoparticles from the walls of the tube. If this background is high relative to the expected EV concentrations, consider a different tube type.

#### 5. EV Analysis

1. For particle size and particle number concentration measurements, we used a nanoflow cytometer (NanoFCM Flow NanoAnalyzer). Techniques such as nanoparticle tracking analysis and resistive pulse sensing can also be used, keeping in mind that each instrument and measurement technique has its own sensitivity range. The NanoFCM, equipped with a 488 nm laser, was calibrated using two types of reference beads: 250 nm silica beads to determine particle concentration and a predefined mixture of silica nanospheres (68, 91, 113, and 155 nm) to establish particle size. PBS was used as a blank control to account for background signal. EV samples (1 µL) were diluted 1:100 or 1:200 in DPBS to achieve an optimal particle count (less than 12,000 particles as specified in the manual). Events were recorded for 1 minute. Particle concentration and size were determined based on flow rate and side-scatter intensity, respectively. From a starting amount of 8 to 12 retinas, we obtained an average of 2.2 x 10^9^ particles/ml/mg retina tissue or 1.2 x 10^10^ particles/ml/retina. The average particle size was 76.3 ± 5.6 nm with a positively skewed distribution (**Figure 2**).
2. For protein analysis, we used the Unchained Labs (Leprechaun; previously ExoView) single-particle interferometric reflectance imaging sensor (SP-IRIS). Silicon chips coated with antibodies against CD81, CD63, and CD9, along with mouse IgG control (Mouse Tetraspanin Chips), were scanned prior to sample addition to establish baseline background signal. EV samples were diluted to 1×10^8^ particles/mL in DPBS to prevent chip saturation. A total of 35 μL of each diluted sample was added to the chips and incubated overnight at room temperature in sealed, light-protected 24-well plates placed on a vibration-free bench. Microscopic images were subsequently acquired using the integrated self-focusing microscope. Retinal EVs demonstrated a predominance of CD9 expression, with lower CD81 expression. CD63 expression was minimal (**Figure 3**). Of note, other single-particle or bulk detection techniques can be substituted. MISEV recommendations also include assessment of depleted markers, specifically proteins enriched in non-EV components. Due to the limited volume of retinal EVs, this could not be performed using Western blotting. Liquid chromatography-mass spectrometry (LCMS) analysis showed no detectable markers of the endoplasmic reticulum (calnexin, GRP94) or Golgi apparatus (GM130). Markers associated with mitochondria (Cytochrome c oxidase subunit 7B) and the nucleus (Histone H2B type 1-F/J/L) were detected, although their peptide-spectrum match counts (# PSMs) were low. In contrast, the cytoskeletal proteins actin and tubulin β4A exhibited higher # PSM values. Retinal ECM components were not present, indicating that ECM components were not enriched.
3. We used transmission electron microscopy to visualize EV morphology (**Figure 4**). Ten µl of sample was thawed and adsorbed to glow-discharged carbon-coated 400-mesh copper grids by flotation for 2 minutes. The grids were washed with three drops of Tris-buffered saline and negatively stained in two consecutive drops of 1% uranyl acetate (UAT) with tylose (1% UAT in deionized water double filtered through a 0.22 μm filter), blotted, then aspirated to cover the sample with a thin layer of stain. Grids were imaged on a Hitachi 7600 TEM operating at 80 kV with an XR80 charge coupled device (8 megapixels, AMT Imaging). Other types of electron microscopy, super-resolution microscopy, and atomic force microscopy can also be used to obtain information about single EVs and their properties.

## Discussion, Potential Pitfalls, and Troubleshooting

Retinal EV studies have primarily relied on cell culture-derived EVs rather than retinal tissue-derived EVs. For example, retinal explants have been cultured *ex vivo* to characterize EV release into supernatant, with polymer-based precipitation used for EV concentration [12]. Cultured retinal pigment epithelium cells are another widely used source of EVs, typically enriched from supernatant following centrifugation [13, 14], and some workflows incorporate density gradients to improve purity [4, 15]. In addition, mouse retinas have been dissociated to generate primary photoreceptor cultures, from which EVs were subsequently collected [16]. While these approaches can reduce contamination from cellular and extracellular matrix debris, they introduce culture-related variables that can affect EV yield and composition [11]. These limitations necessitate robust methods for direct EV separation from retinal tissue to preserve native EV populations and more accurately reflect *in vivo* biology.

**Figure 2.**
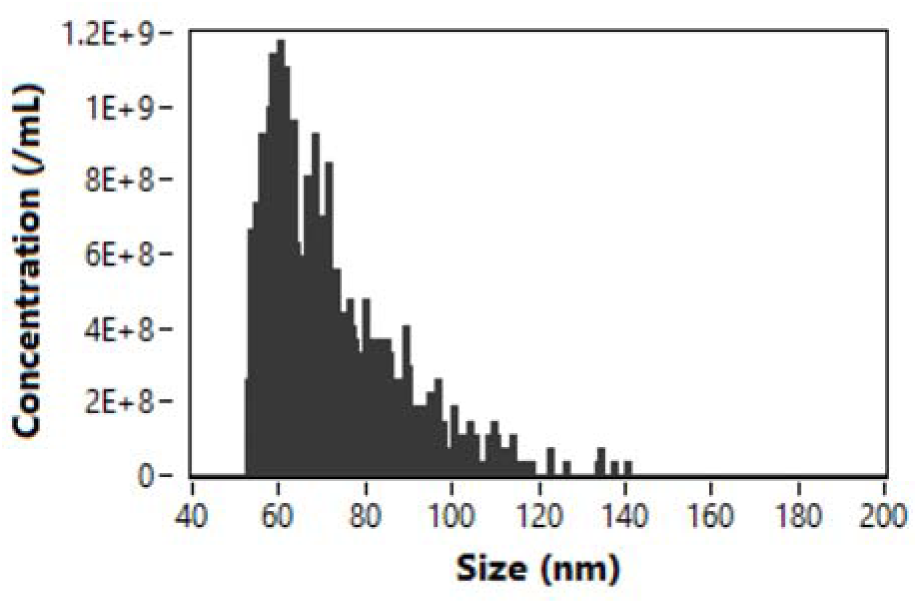
Sample nanoflow cytometer readout showing extracellular vesicle-enriched particle sizes with a positively skewed distribution.

**Figure 3.**
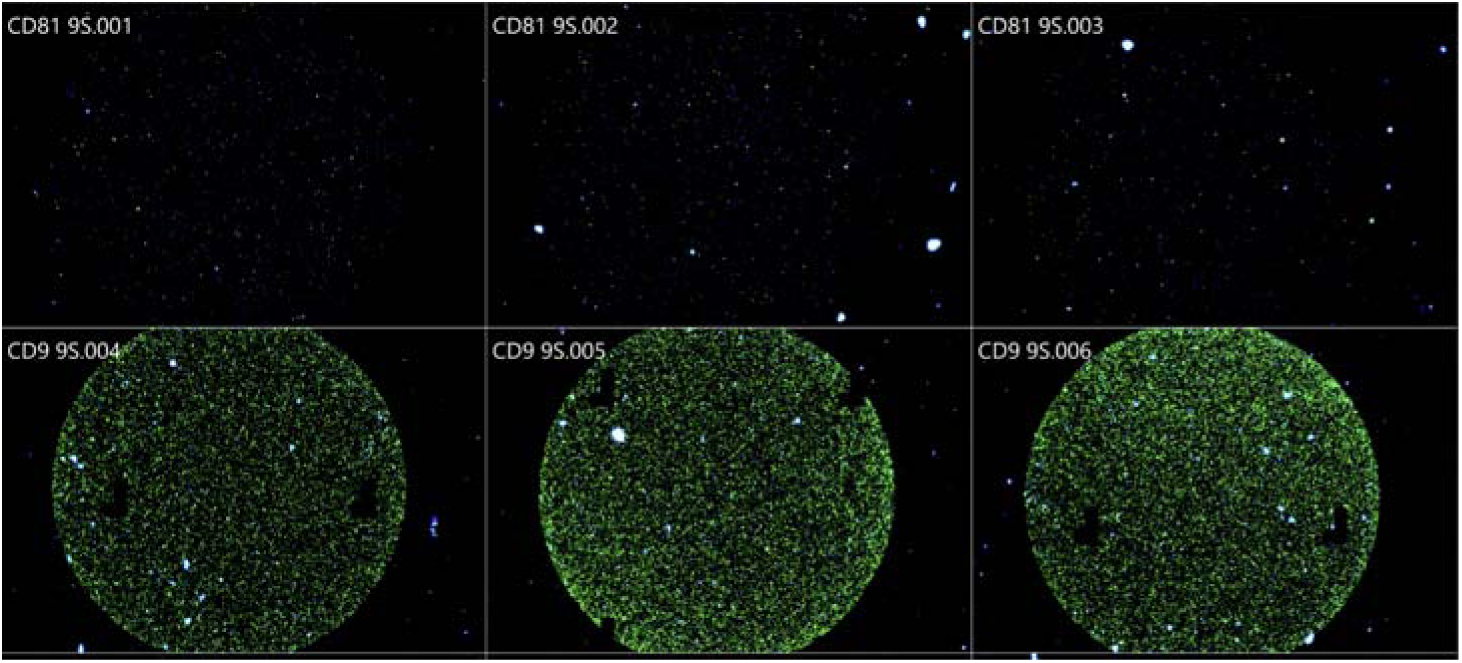
Sample SP-IRIS results showing surface expression of CD81 and CD9 markers of mouse retina extracellular vesicles.

**Figure 4.**
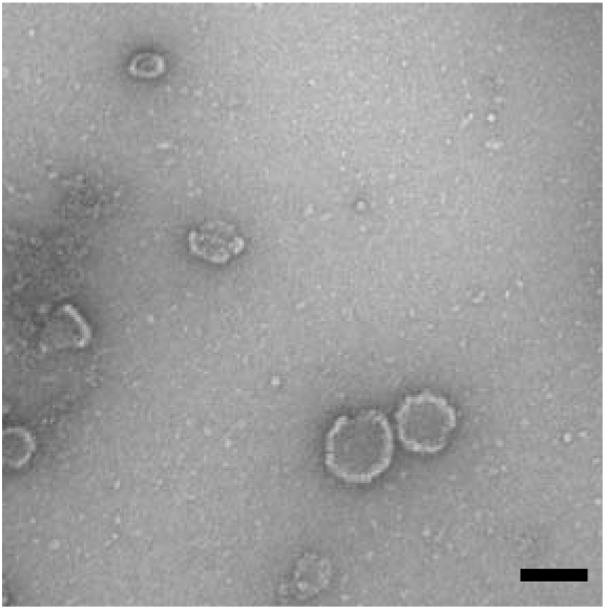
Transmission electron microscopy demonstrating extracellular vesicles (bar = 100 nm)

Direct separation of extracellular vesicles from retinal tissue has been reported but remains comparatively under-developed relative to cell culture-based approaches. In a previously published protocol, retinal tissue was enzymatically digested, centrifuged at low speeds to remove intact cells, and subjected to ultracentrifugation to concentrate vesicle-sized particles [17]. This ultracentrifugation-based approach has been used in subsequent studies [18, 19]. However, ultracentrifugation alone is an enrichment step that can co-isolate non-vesicular protein depending on processing conditions and sample complexity [20].

To address these limitations, our workflow incorporates differential centrifugation at 300 × g and 2,000 × g to sequentially remove contaminants, followed by size exclusion chromatography to obtain EV-enriched, free protein-depleted fractions, and lastly, ultracentrifugation at 110,000 × g to concentrate the EVs. The implementation of size exclusion chromatography enhances EV purity while minimizing co-isolation of non-vesicular material [8]. Additionally, our dissociation uses collagenase rather than papain/DNase. By specifically targeting the extracellular matrix, collagenase is expected to better preserve cell surface markers and maintain membrane integrity. Enzyme choice and exposure time are known to influence EV yield and composition [21, 22].

Several factors can lead to low EV yield or introduce variability in EV characteristics, including contents. Careful optimization and inclusion of appropriate controls are essential.

### Low EV yield

Potential causes of low EV yield include: 1) enzymatic degradation during retina dissection, 2) suboptimal collagenase digestion, 3) loss of pellet or incomplete resuspension following ultracentrifugation, and 4) EV degradation.

Prolonged handling or warm conditions can degrade EVs or alter their composition. Tissue damage may also release intracellular vesicles, confounding downstream analyses. Therefore, we recommend performing dissections rapidly under cold conditions and transferring tissue to dry ice immediately. It is also important to minimize mechanical stress during handling to avoid artificial vesicle release.

Suboptimal collagenase digestion can result from both over-digestion, which can damage cells and generate intracellular vesicle artifacts, and under-digestion, which can result in incomplete tissue dissociation and reduced EV recovery. Therefore, we recommend titrating collagenase concentration and digestion time with a time course to identify optimal conditions. If necessary, a no-enzyme control can help determine the contribution of mechanical dissociation alone.

Small or translucent pellets following ultracentrifugation may be difficult to visualize and can therefore be easily lost during handling. In addition, variability in centrifugation conditions can contribute to inconsistent recovery, and EV aggregation or structural damage may occur if pellets are resuspended too harshly. To avoid these issues, we recommend marking the tube orientation to aid in pellet localization, and using the same rotor type, spin parameters, and fill volumes across experiments. Pellets should be resuspended gently to minimize disruption of EV integrity.

Lastly, we recommend minimizing repeated freeze–thaw cycles prior to downstream analysis or applications since these, among other factors, may contribute to EV loss. The final EV preparation should be aliquoted prior to storage at −80 °C. We also recommend limiting transfer of the EV preparation between tubes to minimize EV loss, as EVs can adhere to tube walls.

### Inconsistent EV characteristics and contents

Potential causes of inconsistent EV characteristics and contents include: 1) blood contamination from incomplete perfusion, 2) intracellular vesicle contamination from tissue damage, and 3) protein contamination. It is also essential to process all samples within the same experimental batch to ensure accurate comparisons among the different EV preparation groups.

Consistent and thorough perfusion should be done across all samples, with liver clearing serving as a reliable proxy for perfusion efficiency. Although it is not possible to completely eliminate blood-derived contaminants or avoid contamination introduced during dissection (such as keratins), these sources of variability should be minimized as much as possible through careful technique and standardized handling procedures. Moreover, comparing perfused versus non-perfused tissues can help quantify the impact of blood contamination on downstream EV analyses.

Intracellular vesicle contamination can arise from tissue damage or harsh processing conditions. To minimize this, gentler dissection techniques should be used, including shorter handling durations, reduced pipetting force, and larger pipette openings to limit shear stress. As noted above, over-digestion during enzymatic treatment can also compromise cell integrity resulting in intracellular vesicle artifacts. Digestion time should therefore be titrated as described above.

Lastly, protein contamination may arise from excessive enzymatic digestion or overly vigorous mixing, both of which can release soluble proteins into the EV-enriched fraction. In addition, overloading SEC columns can reduce separation efficiency, resulting in co-elution of EVs and protein contaminants. SEC loading volumes should be optimized to avoid column saturation. Collecting narrower fractions can improve separation resolution between EVs and soluble proteins. If there are concerns regarding contaminants, we recommend validating the EV-enriched fractions using EV markers (such as CD9 or CD81) to confirm enrichment and purity of EV-containing fractions.

One key limitation is that photoreceptor outer segments are vesicle-rich structures, and membrane fragments derived from these compartments may co-isolate with EVs. This potential confounder should be noted in any study of retinal EVs. Future studies are needed to identify how true EVs differ from other vesicles released from retinal tissue, allowing better interpretation of the contribution of EVs.

In summary, separating EVs from other components of mouse retinal tissues is essential to study the role of EVs *in vivo* physiological and pathological state of retinal cells. Here, we describe an optimized workflow integrating differential centrifugation, size exclusion chromatography, and ultracentrifugation to improve EV recovery from intact retina while preserving EV integrity. In accordance with MISEV guidelines, robust characterization using particle analysis, protein profiling, and imaging-based approaches remains critical for validating EV preparations. Overall, this protocol provides a reproducible framework for isolating retinal EVs directly from tissue, enabling more physiologically relevant downstream molecular and functional studies.

## Acknowledgements

The authors thank Tanina Arab, PhD, for her valuable advice on experimental methods.

